# Autorepression of Yeast Hsp70 co-chaperones by intramolecular interactions involving their J-domains

**DOI:** 10.1101/2024.02.05.578849

**Authors:** Mathieu E. Rebeaud, Satyam Tiwari, Bruno Fauvet, Adelaïde Mohr, Paolo De Los Rios, Pierre Goloubinoff

## Abstract

The Hsp70 chaperones control protein homeostasis in all ATP-containing cellular compartments. J-domain proteins (JDPs) co-evolved with Hsp70s to trigger ATP-hydrolysis and catalytically upload various substrate polypeptides in need to be structurally modified by the chaperone. Here, we measured the protein disaggregation and refolding activities of the main yeast cytosolic Hsp70, Ssa1, in the presence of its most abundant JDPs, Sis1 and Ydj1, and two swap mutants, in which the J-domains have been interchanged. The observed differences by which the four constructs differently cooperate with Ssa1 and cooperate with each other, as well as their observed intrinsic ability to bind misfolded substrates and trigger Ssa1’s ATPase, indicates the presence of yet uncharacterized intra-molecular dynamic interactions between the J-domains and their remaining C-terminal domains. Taken together, the data suggest an auto-regulatory role to these intra-molecular interactions within both type A and B JDPs, which might have evolved to reduce energy-costly ATPase cycles by the Ssa1-4 chaperones that are the most abundant Hsp70s in the yeast cytosol.

**Graphical abstract:**
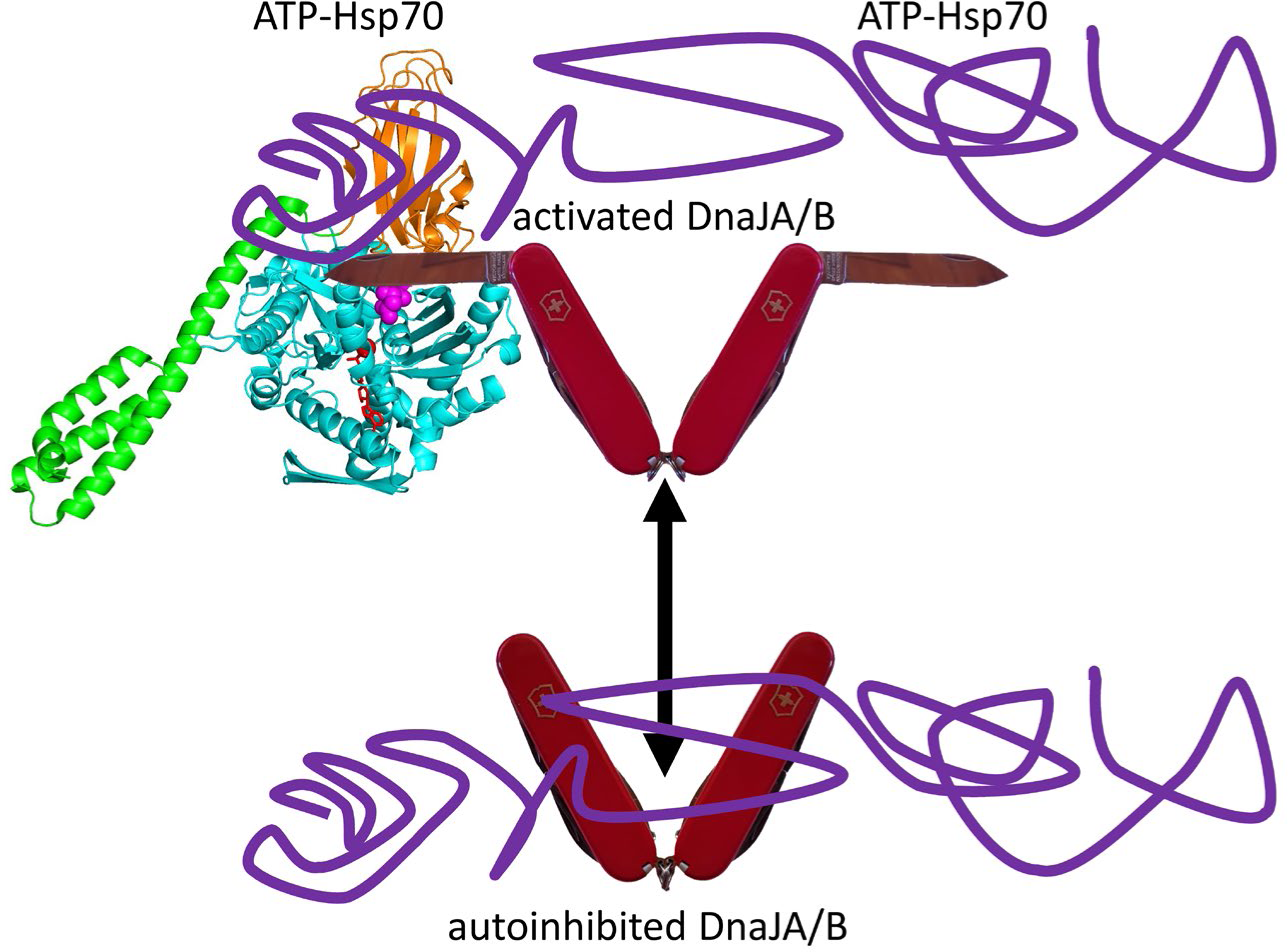
Lower panel: autoinhibited DnaJA or DnaJB dimers, drawn here as Swiss army knives with sequestered J-domains as folded blades, can bind misfolded polypeptides (violet). Upper panel: DnaJA or DnaJB become active when their J-domains are exposed and can bind ATP-Hsp70s, and transfer the misfolded polypeptides, respectively, onto Hsp70’s nucleotide binding (Cyan) and protein binding domains (Orange and Green). Hsp70’s interdomain linker (DLLLLDV, Magenta).

## Introduction

During *de novo* folding and under stress, many native proteins may transiently denature, readily misfold and form stable aggregates lacking their specific dedicated biological activity (Fauvet, Rebeaud et al. 2021). Misfolded species can be cytotoxic and, in humans, are associated with several neurodegenerative disorders, such as Alzheimer’s and Parkinson’s diseases (Sweeney, Park et al. 2017). Early in Life’s history, the first prokaryotes evolved a complex protein quality control network comprising several classes of proteins, including highly conserved molecular chaperones that can specifically target, bind, and thereby repair polypeptides that are conformationally compromised and highly conserved ATP-fueled chaperone-gated endo-cellular proteases that can specifically target, bind and degrade potentially toxic polypeptides who became irreversibly misfolded (Fauvet, Rebeaud et al. 2021, Rebeaud, Mallik et al. 2021). Except for the Hsp20s, the main chaperone families, Hsp100s, Hsp90s, Hsp60s, and Hsp70s are ATPases that can use the energy liberated by ATP hydrolysis to remodel bound misfolded and aggregated protein substrates, ultimately leading to refolding to the native state. Among the ATPase chaperones, the 70 kDa Heat Shock Proteins (Hsp70s) have emerged as the central hub of the protein quality control network that coordinates the optimal unfolding of stably misfolded or alternatively folded protein species, leading to the proper refolding of native proteins, even under non-equilibrium conditions unfavorable to the native state (Goloubinoff, Sassi et al. 2018, Rebeaud, Mallik et al. 2021).

The triage of polypeptide substrates in need to be structurally altered/modified by Hsp70s is initially performed by their obligate J-Domain Protein (JDP) co-chaperones (Kampinga and Craig 2010, Zhang, Malinverni et al. 2022, Marszalek, De Los Rios et al. 2023). JDPs feed Hsp70s with polypeptide substrates and promote the locking of Hsp70s onto the incoming polypeptide substrates that are, but not limited to, misfolded and/or aggregated. To catalyze polypeptide-uploading onto Hsp70 and trigger Hsp70’s ATPase, all JDPs must therefore comprise at least two domains: one very diversified that can directly recognize or be co-localized with specific Hsp70 substrates and one remarkably conserved, the namesake J-domain (JD), that recognizes and binds Hsp70 molecules in their ATP-bound conformation much more strongly than when they are in the ADP-bound conformation (Mayer and Kityk 2015). JDs comprise ∼65 residues, some of them highly conserved, like a characteristic His-Pro-Asp (HPD) motif (Kampinga, Andreasson et al. 2019) known to specifically anchor into a pocket of the Hsp70-ATP complex in which it closely interacts with a folded inter-domain linker between the protein-binding and nucleotide-binding domains of ATP-Hsp70 (see PDB 5NRO). In contrast, in the ADP-Hsp70 conformation, it does not interact with the unfolded inter-domain chaperone linker that became exposed and unstructured (Kityk, Kopp et al. 2018) (see PDB 2KHO). The concomitant interaction of a JD and a polypeptide substrate with Hsp70 greatly accelerates Hsp70’s ATPase cycle (Russell, Karzai et al. 1999, Han and Christen 2004, Kityk, Kopp et al. 2018), resulting in a non-equilibrium enhancement of the affinity of the chaperone for its substrates (De Los Rios and Barducci 2014).

Following a denaturing stress, such as heat shock, conserved homodimeric class A and B JDPs (traditionally called Hsp40s, and in yeast Ydj1 and Sis1, respectively) predominantly bind to misfolded and aggregated proteins (Kampinga and Craig 2010, Zhang, Malinverni et al. 2022) by their two C-terminal domains (CTDs) (including the structurally poorly-resolved linkers with the JDs) and, at the same time, bind ATP-bound Hsp70s through their two N-terminal JDs. Owing to the high flexibility of G/F-rich linkers connecting the JDs to the CTDs, the JDs of both classes would be expected to freely swing and, thereby, seek unrestrained binding to ATP-Hsp70s. Yet, there is emerging evidence of intramolecular regulatory mechanisms that may reversibly block the interaction between the JDs of class B JDPs (yeast Sis1 and human DNAJB1) and the corresponding cytoplasmic Hsp70s (yeast Ssa1-4 and human HSC70). In the case of Sis1, interactions between Glu-50 on JD and Arg-73 are documented on a short helical motif immediately C-terminal to helix IV (Yu, Ziegelhoffer et al. 2015) (Fig.S1), whose mutational disruption, or by swapping the JD of Sis1 for that of Ydj1, increases the ability of Sis1 to trigger the refolding activity of Ssa1 (Yan and Craig 1999, Yu, Ziegelhoffer et al. 2015). It has also been reported that human DNAJB1 comprises a distal α-helix inside the G/F region (helix V, Fig.S1 and 2, sequences in Supplementary Information), whose deletion facilitates the cooperation of DNAJB1 with HSC70 for refolding different misfolded substrates (Fan, Lee et al. 2004, Faust, Abayev-Avraham et al. 2020). The CTDs of DNAJB1 also participate in this intricate network of interactions, as testified by the finding that the C-terminal EEVD motif of HSC70, which interacts with the CTD1 subdomain of DNAJB1 (Jiang, Rossi et al. 2019, Johnson, Nadel et al. 2022), is necessary to alleviate the helix V-induced inhibition of DNAJB1’s co-chaperone activity, however through a mechanism yet not fully resolved structurally. Similar results have been obtained for yeast Sis1 (Wyszkowski, Janta et al. 2021). Furthermore, through a series of modular domain deletions or domain swaps with Ydj1, it has been shown in yeast that only chimeras containing the CTDs of Sis1 are effective in preventing prion propagation and toxicity, with their effectiveness depending to some extent on the origin of the JD and of the G/F region (Kumar, Reidy et al. 2021). In contrast to Sis1, the presence of an intramolecular mechanism in Ydj1 that might tune the action of its JD has not been reported, despite AlphaFold2 (Jumper, Evans et al. 2021) predicting inter-domain motifs similar to those of Sis1: two alfa-helices, one right following helix IV and another, although with lower reliability, within the G/F region (see pLDDT in Fig.S3) (Wall, Zylicz et al. 1995, Yan and Craig 1999, Cajo, Horne et al. 2006). In addition to these examples, recent experimental studies have documented the interaction of the JD with other structural elements of the same polypeptide in various JDPs (*e.g.* DNAJB6, DNAJB8) (Karamanos, Tugarinov et al. 2019, Ryder, Matlahov et al. 2021). Adding to the richness of effects that are modulated by interactions between the JDs and other domains of JDPs, it was found that the ability of individual DNAJA or DNAJB homodimers to unlock the unfolding-refolding activity of Hsp70s is much higher when they are in the presence of one another, as compared to their stand-alone effects (Nillegoda, Kirstein et al. 2015).

In this work, we investigated the presence of intramolecular mechanisms by which the two main JPDs in the yeast cytosol, Ydj1 (class A) and Sis1 (class B) (hereafter called YY and SS respectively), regulate the activity of the main yeast cytosolic Hsp70, Ssa1. Inspired by previous works (Yan and Craig 1999, Johnson and Craig 2001, Fan, Lee et al. 2004, Yu, Ziegelhoffer et al. 2015, Schilke, Ciesielski et al. 2017), we developed two new chimeras, henceforth called YS and SY, obtained by swapping the J-domains of YY and SS (with the first letter indicating the parent protein of the N-terminal JD and the second letter indicating the parent protein of the remaining part of the chimera sequence). Comparing the behavior of the two wild-type JDPs with that of the two chimeras across a series of assays (disaggregation and refolding of pre-aggregated enzymes, prevention of aggregation, and activation of Ssa1’s ATPase), we highlighted the presence of intramolecular interactions between the JDs, the G/F regions and the CTDs that likely offer cells a way to regulate their multiple Hsp70- and JDP-dependent functions, and thus the energy-consuming cycles by their abundant Hsp70s, depending on cellular needs.

## Results

We designed the two chimeras by swapping the 80 N-terminal residues containing the JD domains and the first 10-12 residues of the G/F hinge region of Ydj1 and Sis1, (Fig.1, Supplementary Information for the detailed sequences). In the case of Sis1, the swap preserved the inhibiting E50-R73 interaction but abolished the interaction between the JD and helix V.

**Figure 1:**
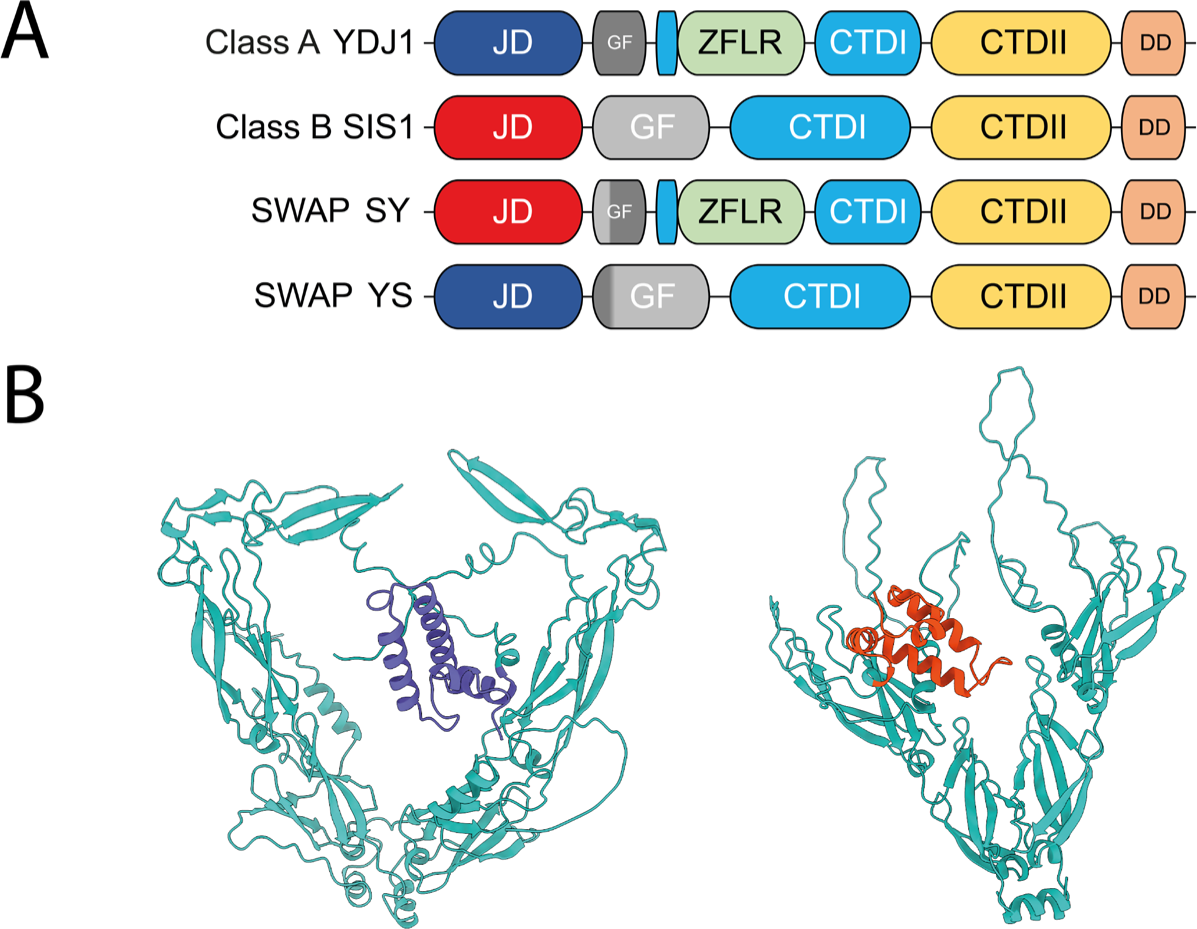
Domain organization of yeast class A (YDJ1) and B (SIS1) JDPs and of the swap chimeras. A: Domain organization of Sis1 and Ydj1. J-domain (JD), G/F rich-region (GF), Dimerization Domain (DD), Zinc-finger-like region (ZFLR). B: Alphafold2 prediction of left: Class A (YY) and right: Class B (SS) with only one J-domain of each homodimer shown (complete model in Fig.S2). For more clarity, the model of SS has been rotated 90° on the left compared to the one of YY.

We then tested the *in-vitro* ability of increasing concentrations of YY, SS, YS, and SY to drive the native refolding of stable, preformed urea- and heat-aggregated luciferase (Shorter 2011) by a constant amount of Ssa1, Sse1 (the yeast member of the Hsp110 family), which in the presence of ATP is considered to act both as a Nucleotide Exchange Factor (NEF) and as an ATP-dependent disaggregating co-chaperone (Fig.2). Remarkably, at all concentrations, YS was found to be much more effective at refolding pre-aggregated luciferase than wild-type SS, with a more marked difference at low concentrations. Similarly, SY was more effective than YY, although both were much less effective than SS. The action on different substrates, preformed heat-aggregated G6PDH and MLucV (Tiwari, Fauvet et al. 2023), showed a similar pattern (Fig.S4 and 5).

**Figure 2:**
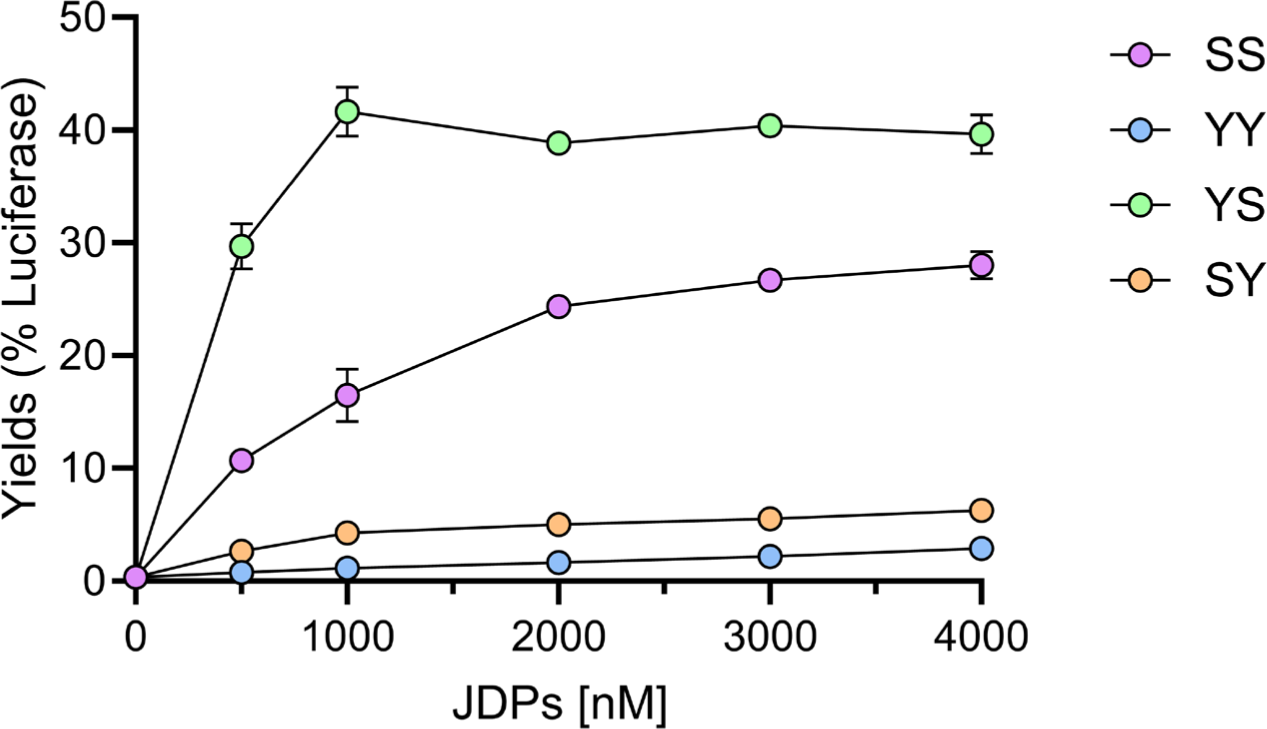
Different refolding effectiveness of wild-type and chimeric JDPs. Efficiency of different JDPs proteins (SIS1 and YDJ1 and the two swap SY and YS) to power SSA1-mediated disaggregation of heat-urea pre-aggregated Luciferase. 0.2 µM of stable inactive luciferase aggregates were incubated for up to 120 minutes at 25°C in the presence of 5 mM ATP, 6 µM of SSA1, 0.75 µM SSE1, and between 0 to 4 µM of either YY, SS, SY or YS. In all panels, error bars represent mean ± SD (n = 3).

To explain the observed ranking of effectiveness, it was necessary to invoke some functional interaction between the JDs and the CTDs. Indeed, the independent-domain assumption could explain the greater yield of YS compared to SS by positing that the JD of YY was more effective than the JD of SS at recruiting and activating Hsp70, while the CTD of SS was more effective than the CTD of YY at capturing the substrate and delivering it to Hsp70. Yet, a direct extension of this argument would have predicted that SY, which would have comprised the least effective instance of each domain, should have been the least efficient of the four constructs, contrary to our findings. The independent action hypothesis can thus be excluded, and intramolecular interactions between the JDs and the CTDs were required to explain the results.

We thus addressed more in detail the effects of the JD swap on the two functions usually ascribed to the two domains: stimulation of the ATPase cycle of Ssa1 by the JDs and substrate binding by the CTDs (including most of the G/F hinge and, in the case of Ydj1, also the cysteine-rich domain).

To test substrate-binding by SS, YY, YS and SY, we measured their specific ability to block the increase of the light-scattering signal of aggregating, urea-pre-unfolded luciferase (see Methods, and Fig.S6 and 7 for a characteristic time-course of aggregation). Consistent with the literature, different doses of YY were found to be more effective at reducing protein aggregation than SS (Lu and Cyr 1998, Yan and Craig 1999). Comparing the swap-mutants with the wild-types, we observed, in several assays and for different aggregating substrates, that SY systematically prevented aggregation more efficiently than YY (Fig.3 and Fig.S7 and 8). These results offered insights on the intramolecular interactions of these JDPs. Barring in mind the unlikely possibility that the JD of YY can by itself bind substrates, a feature that has never been reported, the differences found between YY and SY suggested that strong intra-molecular interactions between the JD of YY and its CTD partially interfered with the ability of the YY CTD to engage aggregating polypeptide substrates. In contrast, in the SY chimera, the JD of SS has not co-evolved, and therefore is not expected to have transient interactions with YY’s CTDs, allowing them to freely bind misfolding substrates and prevent their aggregation. In contradistinction, the reverse chimera, YS, was only marginally, if at all, worse than SS at preventing substrate aggregation (see Fig.S8 for additional substrate), indicating that any intra-SS JD/CTD interactions, if present, do not interfere with substrate binding.

**Figure 3:**
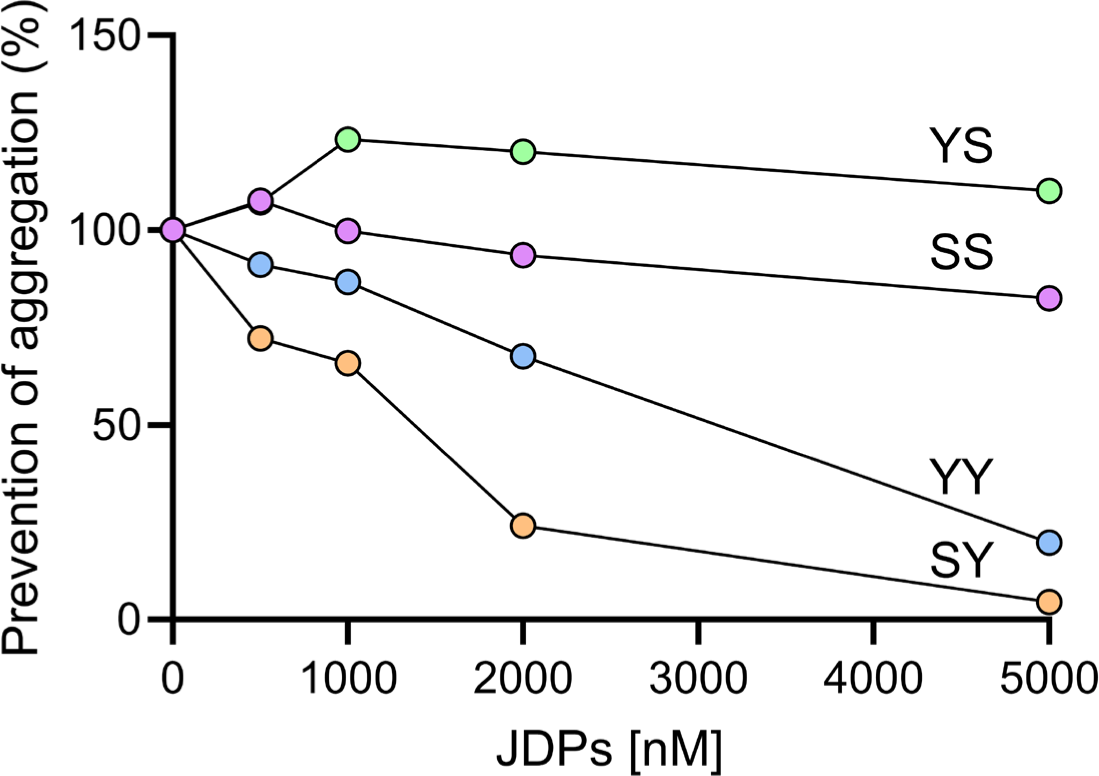
J-domain protein acting as holding chaperone. Efficiency of SS, YY, SY, and YS at preventing luciferase aggregation. 10 µM Luciferase was pre-unfolded with 6 M urea at 30℃ for 30 min, then diluted to a final concentration of 0.3 µM in buffer (50mM Tris-HCl pH 7.5, 150mM KCl, 10mM MgCl2, 2mM DTT), in the absence or in the presence of increasing concentrations of YY, SS, SY, YS. Relative aggregation yields following 800 seconds were calculated in the presence of increasing concentrations of YY, SS, SY, YS. Maximal aggregation without JDPs at 800 seconds was set to 100%. the time-dependent aggregation was monitored by light scattering at 340 nm at 30 ℃.

To test the second relevant role of JDPs, namely stimulation of the ATPase cycle of Hsp70, we measured the rates of ATP hydrolysis by Ssa1 in the presence of increasing concentrations of the four JDPs (Fig.4A) and, as a reference, we also measured the ATPase rates in the presence of the same increasing concentrations (protomer by protomer) of isolated JDs.

**Figure 4:**
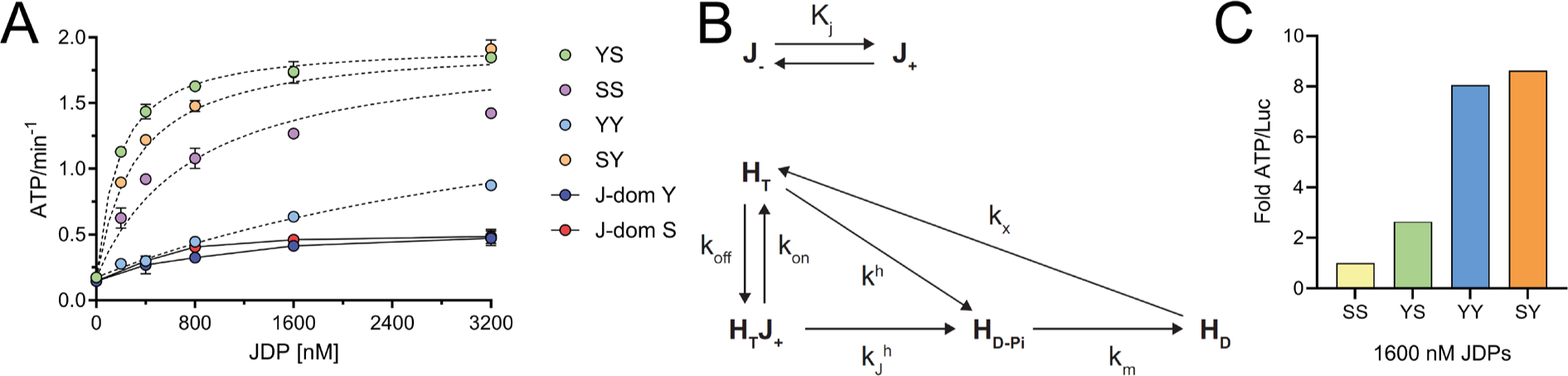
ATPase activity of Ydj1, Sis1 and of their J-domain swapped chimeras. A) Rates of ATP hydrolysis of 6000 nM SSA1 and 750 nM SSE1 without substrate in the presence of increasing concentrations of YY, SS, YS, and SY. B) Reaction scheme for the activation of the ATPase cycle of Ssa1 by the different Hsp40 constructs. C) Fold-change of ATP hydrolyzed per natively refolded luciferase with SIS1 set as a reference, by 6000 nM SSA1 and 750 nM SSE1, in the presence of either 1600 nM YY, SS, SY, or YS. The error bars in panels A represent mean ± SD (n = 3).

No major differences were found in the effects of the isolated JDs of YY and SS, indicating that their ability to interact with Ssa1 and to stimulate its ATPase activity is similarly low (Fig.4A). The effects of the two swap chimeras, YS and SY, were also similar, albeit a much higher ATPase stimulation in their presence compared to the isolated JDs, suggesting that some parts of the full-length JDPs different from the JDs might also act as Ssa1 substrates (Laufen, Mayer et al. 1999, Kityk, Kopp et al. 2018), thereby synergistically stimulating Ssa1’s ATPase. The wild-types YY and SS had intermediate effects, between those of isolated JDs and those of the swap chimeras. SS was systematically less effective than both chimeras, indicating that its JD was also in an inactive state that affected the ability of SS to activate Ssa1. Instead, YY was more effective than isolated JDs only at the highest tested concentrations, and was always much less effective than SS. Since the JDs of Ydj1 and Sis1 were similarly effective both when isolated and when swapped, the decreased effectiveness of wild-type YY and SS suggested that intramolecular interactions between their respective JDs and the CTDs were inhibiting their ability to freely interact and activate Ssa1’s ATPase. This auto-repression was particularly strong in the case of YY.

We modeled these data by means of the reaction schemes in Fig.4B. YY and SS are in a dynamical equilibrium between the auto-repressed and active states (J_-_ and J_+_ respectively), characterized by an equilibrium constant K_J_ , associated to the relation [J_+_] = K_J_ [J_-_] (red reaction): K_J_ values smaller than 1 mean that the JDP is most likely found auto-repressed. Ssa1 can only bind to J_+_, with JDP association and dissociation rates k_on_ and k_off_. Following JDP association, ATP hydrolysis is induced with rate k_h_^J^, with the ensuing release of inorganic phosphate. This reaction is followed by ADP-to-ATP exchange with a rate k_x_. In parallel to the JDP induced hydrolysis pathway, Ssa1 can also hydrolyze ATP spontaneously, albeit at a much slower rate. We simplified the scheme for the swap chimeras with the assumption that they are never auto-repressed (due to the presumed absence of complementary surfaces on their JD and CTD, that in wild type SS and YY would have co-evolved to specifically interact to form the auto-repressed state), and that thus no J_-_ state is present. The analytical solution of these reactions provides formulas for the steady-state ATPase rate that we used to fit the experimental data (Fig.4A, dashed lines). While the fits were not perfect, due to some simplifications in the model and in the potential presence of unknown features of the system that we were not capturing, they were nonetheless instructive. The main result of this analysis was that in both wild-types, the equilibrium was tilted toward the auto-repressed state (K_J_ < 1), much more strongly in YY (K_J_≈1/30) than in SS (K_J_≈1/2). This result further supported the hypothesis that intramolecular interactions in YY and SS, involving the JD, may partially sequester the JD and/or the parts of the JDP that could act as Ssa1 substrates.

An ancillary observation from Fig.4A is that in all cases, apart from YY, saturation of the Ssa1 ATPase activity was achieved by amounts of JDP or of isolated JDs that were sub-stoichiometric to Ssa1. Within our reaction scheme (Fig.4B), this is a consequence of the overall exchange rate (namely, release of Pi and ADP-to-ATP exchange) that was much slower than the rate of ATP hydrolysis and of the ensuing release of the JD. This implies that, in the absence of NEF, a single JD acts catalytically, activating several Ssa1s before the first one undergoes nucleotide exchange and resets its cycle (Kityk, Kopp et al. 2018). This is consistent with proteomic data estimating the presence in the yeast cytosol of seven times more Ssa1-4 (together ∼14 µM protomers) than of YY and SS (together ∼2 µM protomers) (Ghaemmaghami, Huh et al. 2003, Lawless, Holman et al. 2016, Mackenzie, Lawless et al. 2016).

The addition of a true substrate (pre-aggregated luciferase) further increased all ATPase rates 1.5-1.7-fold for all full-length JDP constructs (Fig.S9). Combining these data with those shown in Fig.2A, we estimated the relative ATP cost of the *in vitro* disaggregation/refolding reaction in the presence of a constant amount (1600 nM) of SS, YY, YS or SY (Fig.4C). The most energy efficient JDP of the four constructs was SS (whose relative efficiency was set to 1). The second most energy-efficient JDP was YS, which nevertheless hydrolyzed 2.6-times more ATPs per refolded Luciferase, while YY and SY were equally energy inefficient, hydrolyzing at least 8 times more ATP per refolded luciferase than SS. Thus, the constructs with the CTDs of Ydj1 were both slower at refolding pre-aggregated luciferase, and less efficient at using ATP energy.

In metazoa, as well as in yeast, the disaggregation and subsequent refolding of luciferase aggregates is enhanced by the synergistic co-action of homodimeric class A and B JDPs, likely mediated by intermolecular interactions between JDs and CTDs of different classes (Nillegoda, Kirstein et al. 2015, Wyszkowski, Janta et al. 2021). We first confirmed these results, finding that in our assays a mixture of SS and YY at constant total JDP concentration was more effective than the sum of their individual contributions (Fig.5A). In particular, we observed that the YS chimera and the YY:SS mixture had was very similar behavior for equal YS and SS concentrations. These data suggested that the mutual action of SS and YY is catalytic: small amounts of each can greatly enhance the activity of the other, with the overall effect matching the one of YS, the most active chimera. We confirmed that interpretation by measuring the effects of increasing concentrations of YY (SS) on fixed amounts of SS (YY) (Figs.5B and C, Fig.S10A). In both cases, small amounts of one JDP were able to enhance the activity of the other. In order to assess the difference in activation between the YY:SS and YS:SY pairs, we also compared their activatory effect by adding an increasing quantity of YS to a decreasing quantity of SY, with a fixed quantity of 4000 nM total. The pair of wildtype JDPs showed an enhanced activatory effect in the mixture, whereas the pair of SWAPs were no more activated than the theoretical value of the mixture (Fig.S10A for the wild-type mixture and B for the mixture of swap chimeras).

**Figure 5:**
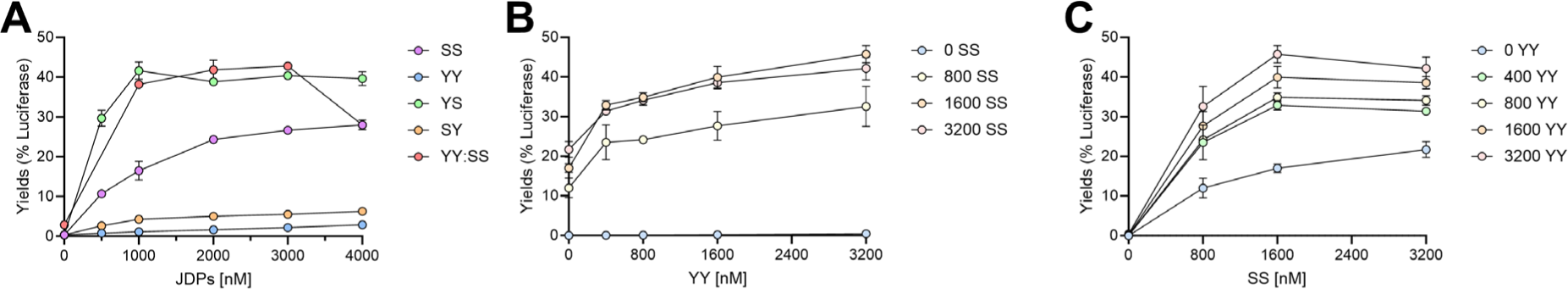
Synergistic action of Ydj1 and Sis1. A) Data as in Fig.2, together with data for the measured yields of refolded luciferase achieved by a x%/(100-x)% mixture of SS and YY respectively. The simple sum of the two contribution is in dashed line. B) Dose response of YY for fixed amounts of SS to refold pre-aggregated luciferase. C) Dose response of SS for fixed amounts of YY to refold pre-aggregated luciferase. The error bars in panels B and C represent mean ± SD (n = 3).

## Discussion

There is emerging evidence that intramolecular dynamic interactions in some JDPs might have a role in reducing unnecessary interactions with Hsp70s, or example when there is no stress and very little protein aggregation, and thereby possibly avoiding futile ATP hydrolysis in unstressed cells, by maintaining sequestered, at least partially, their J domains. This is evocative of a Swiss army knife (see graphical abstract), whose blade can either be autorepressed, with its blade folded in its handle, or derepressed with its blade exposed and poised to perform its cutting function. Our data suggest that both DnaJAs and DnaJBs can either be auto-repressed, with their JDs strongly interacting with their C-terminal domains, or derepressed with their JDs exposed and free to activate Hsp70. Whereas the purpose of knife’s auto-repression is security, it is tempting to speculate that JDPs autorepression may reduce unnecessary futile ATP hydrolysis by the Hsp70s, which are abundant not only in stressed cells, but also in quiescent unstressed yeast cells deprived of energy sources. In the case of human DNAJB1, a class B JDP implicated in protein refolding and disaggregation, autorepression has been ascribed to two intramolecular structural elements: a contact between E50 and K73 (E50 and R73 in yeast Sis1, relevant for the present work) and a more extensive interaction, which was also observed experimentally and predicted by AlphaFold 2, between the JD and a helix (helix V) in the G/F region (Fig.S1). Noticeably, AlphaFold 2 also predicted strong interactions between Sis1’s JD and a further distal helix in the G/F hinge region (Fig.S1, red helix), which is expected to sterically clash with the activatory interaction of the HDP motif in the JD (Fig.S1 green), with ATP-Hsp70 in the pocket containing the conserved linker (black) connecting protein-binding to nucleotide-binding domains (PDB: 5NRO). Interestingly, binding of the EEVD tetrapeptide at the C-terminal end of Ssa1 to the C-terminal domain of Sis1 also contributes to the abolition of its autoinhibition (Ghaemmaghami, Huh et al. 2003, Lawless, Holman et al. 2016, Mackenzie, Lawless et al. 2016) . A similar activatory effect can also be achieved by specific mutations (*e.g.* E50A, or F102A in helix V), or the full deletion of helix V (Faust, Abayev-Avraham et al. 2020). In yeast Sis1, the presence and role of helix V has not been experimentally determined, albeit its presence is also predicted by AlphaFold 2 (Fig.S3 and AlphaFold resolved structure on Uniprot (UniProt 2015) ID of SIS1: AF-P25294-F1), with contacts with the JD similar to those for DNAJB1. No similar auto-repressed interactions have been documented so far for class A JDPs.

In this work we have used domain-swap mutagenesis, biochemical assays, ATP hydrolysis measurements and chaperone mediated disaggregation and refolding assays and chaperone-binding assays (light scattering and FRET measures) to highlight the presence of auto-repressive interactions between the J-domains and the other C-terminal domains of the two main class A and B JDPs in yeast, Ydj1 and Sis1 respectively. We assessed their effectiveness at performing the two individual functions typical of these JDPs, namely the binding of misfolding and already stably aggregated protein substrates, and the stimulation of Ssa1’s ATPase activity (the most abundant cytosolic Hsp70 in yeast) and compared them with two JDP chimeras in which the J-domains of Sis1 and Ydj1 were interchanged. We found that the SY swap-mutant was more effective than YY at preventing the aggregation of luciferase at all tested concentrations. Since J-domains are not known to be involved in substrate binding, a task that is a prerogative of both the G/F hinge region and of the CTDs, this result hinted at the presence of a specific intramolecular interaction between the J-domains and the downstream domains containing the G/F hinge the cysteine rich domain and the CTD of Ydj1, that would reduce access of the substrate to its binding sites. Exchanging the Ydj1 J-domain for that of Sis1 would then abolish this interaction, allowing unhindered substrate and Hsp70 binding. It is thus tempting to hypothesize that substrates might take part in the relief of the auto-repression, a possibility that should deserve future investigation.

Noticeably, although SS and YS did not have any observable “holdase” activity (Lu and Cyr 1998), both were the most effective JDPs at promoting the unfoldase/refoldase activity of Ssa1. In contrast, YY and SY, which were found to have a strong binding affinity for misfolding and aggregating substrates, were the least Hsp70-activating JDPs in terms of ATP-fueled unfoldase activity, suggesting that the ability of a JDP co-chaperone to bind and prevent protein misfolding aggregation does not directly correlate with its ability to transiently bind and then <u>dissociate</u> and thereby transfer a misfolded polypeptide substrate to Hsp70 and to activate its ATP-fueled unfoldase/refoldase activity.

The results for the stimulation of the ATPase cycle of Ssa1 were in line with these observations. The isolated, and thus unrepressed J-domains of Ydj1 and Sis1 were almost identically poorly efficient stimulators. Likewise, the two swap chimeras were similarly effective, albeit much more than the J-domains alone, suggesting that, as previously reported for bacterial DnaJ, some parts of these JDPs might also act as Hsp70 substrates and synergize with the J-domain in the acceleration of the chaperone ATPase cycle. The observation that the wild-type JDPs had effects intermediate between the isolated J-domains and the chimeras hinted at intramolecular interactions involving the J-domains that lead to their partial inhibition and/or to the sequestration of the regions that would act as Ssa1 substrates, thus partially reducing their ability to stimulate of the ATPase of Ssa1. In particular, the very mild ATPase stimulation by Ydj1, which was detectable only at the highest concentration, despite its J-domain being normally active when alone or paired with the CTDs of Sis1, suggested that the J-domain of Ydj1 was more strongly auto-repressed by intramolecular contacts than the J-domain of Sis1, as our computational model also determined, calling for future investigations to elucidate the structural basis of these hitherto uncharacterized intramolecular interactions.

Taken together, all these observations conjure at explaining the results of luciferase disaggregation and reactivation. YY and SY were the least effective JDPs of the four constructs, either because they were inefficient at binding to large aggregates and to lead to their disaggregation by Ssa1, as proposed elsewhere for human DNAJA1 (Nachman, Wentink et al. 2020), or because of their strong propensity at binding and not releasing unfolded/misfolded proteins, thus interfering with the later stages of the disaggregation/reactivation process (Veinger, Diamant et al. 1998, Moran Luengo, Kityk et al. 2018) . For protein disaggregation by Ssa1 and NEFs of the Hsp110 family, Class B JDPs are known to be more efficient than class A (Nillegoda, Kirstein et al. 2015). Our results confirmed this observation, which is possibly due to the ability of the EEVD C-terminal tetrapeptide of Ssa1 to disengage the J-domain of Sis1 from its auto-inhibitory intramolecular interactions.

Why is the YS chimera the most effective of them all? YS mimicked the behavior of the YY/SS mixture at concentrations similar to those of Sis1. Given the apparent absence of J-domain autorepression in YS molecules, it is tempting to speculate that this is indeed the effect that the YY/SS mixture achieves, by molecular interactions that, while most likely based on J-domain/CTD interactions, have not been yet characterized in detail. It is noteworthy that the co-chaperoning activity of 3200 nM SS is strongly increased by a 16-time smaller amount of YY (which has no activity on its own), evocative of a catalytic process (Fig 5B).

But why would have nature evolved JDPs that are less efficient than what would be possible? While we lack a comprehensive perspective of all optimality criteria that lead evolution to specific solutions, the greater energy efficiency of Sis1 suggests that energy might have been a driving factor, by reducing unnecessary interactions of DNAJAs and DNAJBs with the highly abundant cytosolic Hsp70s, especially when energy sources are scarce. More research will be needed to uncover all the disparate mechanisms and conditions that control this “stop-and-start mechanism” of JDPs.

## Material and Methods

### Strains and plasmids

Wild-type *SSA1*, *SSE1*, *SIS1*, *YDJ1, and* swap chimeras were cloned in the pET-SUMO vector for propagation in *E. coli.* The swap chimeras were constructed by PCR by amplifying the 80 first amino acids (J-domain and a small part of the Glycine-Rich region) with the following primers: JYDJ1-F 5’- GCGAACAGATTGGAGGTATGGTTAAAGAA-3’, JYDJ1-R 5’-GCCAGGACC ACCAGGAGCGCCACCAGCAC-3’, JSIS1-F 5’-GCGAACAGATTGGAGGTATGGTCAAGGAG-3’, JSIS1-R 5’- ACCTGGGAATCCGCCACCAAAGCTTGGACCAC-3’) of the two JDPs and insert them inside the vector of the other one using these set of primers (YDJ1V-F 5’-GGCGGATTCCCAGGTGGTGGATTC-3’, YDJ1V-R 5’- ACCTCCAATCTGTTCGCGGTGAGC-3’, SIS1VF 5’-TGGTCCTGGTGGTCCTGGCGGTGC-3’, SIS1VR 5’- CCAATCTGTTCGCGGTGAGCCTCA-3’) by Gibson Assembly (NEB). All constructs were confirmed by sequencing.

### Purification of proteins

For purification of the His10-SUMO tagged wild-type SSE1, SSA1, SIS1, YDJ1, and swap chimeras, were expressed and purified from *E. coli* BL21-CodonPlus (DE3)-RIPL cells with IPTG induction (final 0.5 mM for SSA1 and SSE1 and 0.2 for YDJ1, SIS1, and swap chimeras) at 18 °C, overnight. Briefly, cells were grown in LB medium + ampicillin at 37 °C to OD600 ∼0.4-0.5. Protein expression was induced by the addition of 0.5 mM IPTG for 3 hours. Cells were harvested, washed with chilled PBS, and resuspended in buffer A (20 mM Tris-HCl pH 7.5, 150 mM KCl, 5% glycerol, 2mM DTT, 20mM MgCl_2_) containing 5 mM imidazole, 1mg/ml Lysozyme, 1mM PMSF for 1 h. Cells were lysed by sonication. After high-speed centrifugation (16000 rpm, 30 min/4°C), the supernatant was loaded onto a gravity flow-based Ni-NTA metal affinity column (2 ml beads, complete His-Tag Purification Resin from Merck), equilibrated and washed with 10 column volumes of buffer A containing 5 mM imidazole. After several washes with high salt buffer A (+150 mM KCl, 20mM Imidazole and 5 mM ATP), N-terminal His10-SUMO (small ubiquitin-related modifier) Smt3 tag was cleaved with Ulp1 protease (2mg/ml, 300 μl, added to beads with buffer (20 mM Tris-HCl pH 7.5, 150mM KCl, 10mM MgCl2, 5% glycerol, 2mM DTT). Digestion of His10 Smt3 was performed on the Ni-NTA resin by, His6-Ulp1 protease. Because of dual His tags, His6-Ulp1 and His10-SUMO display a high affinity for Ni-NTA resin and remain bound to it during cleavage reaction. After overnight digestion at 4°C, the unbound fraction is collected (which contains only the native proteins). Proteins were further purified by concentrating to ∼3mg/ml and applying to a size exclusion column (Superdex-200 increase, 10/30 GE Healthcare) equilibrated in buffer A containing 5 mM ATP. Pure fractions were pooled, concentrated by ultrafiltration using Amicon Ultra (Millipore), aliquoted, and stored at -80 °C. All protein concentrations were determined spectrophotometrically at 562 nm using BCA Protein Assay Kit− Reducing Agent Compatible (cat no. 23250).

The purified proteins were collected, concentrated, and stored at -80°C for further use.

### Luciferase refolding assay

Luciferase activity was measured as described previously (Bischofberger, Han et al. 2003, Sharma, De los Rios et al. 2010). In the presence of oxygen, luciferase catalyzes the conversion of D-luciferin and ATP into oxyluciferin, CO_2_, AMP, PPi, and hν. Generated photons were counted with a Victor Light 1420 Luminescence Counter from Perkin-Elmer (Turku, Finland) in a 96-well microtiter plate format.

### G6PDH refolding assay

Heat-pre-aggregated G6PDH was refolded by the SSA1 chaperone system as described previously for the DnaK chaperone system (Mattoo, Farina Henriquez Cuendet et al. 2014), with the following modifications; 500 nM heat-aggregated G6PDH (final concentration) was reactivated in the presence of 6 μM SSA1, incrementing (0–1 μM) JDPs, 0.75 μM SSE1 (the full SSA1 chaperone system) and 5 mM ATP. G6PDH activity was measured at different times of chaperone-mediated refolding reaction at 25°C.

### ATPase assay (Malachite Green)

Colorimetric determination of Pi produced by ATP hydrolysis was performed using the Malachite Green Assay Kit (Sigma-Aldrich, Switzerland) and as described previously (Lee, Roh et al. 2019). Several concentrations of Hsp70 (SSA1) and JDPs (YY, SS, SY, YS) were mixed with or without substrate (200 nM of pre-aggregated luciferase) and with 1 mM of ATP and incubated for 1 hour at 25°C. 4 µL of each sample was taken and put inside a 96-Well plate with 76 µL of H_2_O. A 20-µl volume of Malachite Green reaction buffer was added, and the samples were mixed thoroughly and incubated at 25°C for 30 min before measuring at 620nm on a plate reader (HIDEX-Sense 425-301, Finland). The rate of intrinsic ATP hydrolysis was deduced by subtracting the signal from ATP in the absence of a chaperone.

### Light Scattering

To monitor the aggregation propensity of urea denatured Luciferase as described previously (Tiwari, Fauvet et al. 2023), 10 μM Luciferase was denatured with 6M urea at 30°C for 10 min, then diluted to a final concentration of 0.3 μM in buffer A (50 mM Tris-HCl pH7.5, 150 mM KCl, 10 mM MgCl2, 2 mM DTT), immediately aggregation was monitored by light scattering at 340 nm at 30°C using Perkin Elmer Fluorescence Spectrophotometer.

### FRET measurements and FRET efficiency calculation

All ensemble relative FRET efficiencies were calculated from maximum fluorescence emission intensities of the donor (ED) and acceptor (EA) fluorophore by exiting the donor only at 405 nm wavelength (Fritz, Letzelter et al. 2013, Wood, Ormsby et al. 2018). Fluorescence emission spectra analysis of MLucV reporter was performed on PerkinElmer LS55 fluorometer. Emission spectra were recorded from 480 to 580 nm wavelength with excitation slit 5 nm and emission slit 10nm. Average intensity values of spectral crosstalk were minimized by excitation donor at 405nm. Samples acquired in the same conditions. The relative FRET efficiencies were calculated using the following equation:

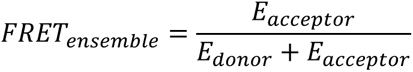

Normalized FRET efficiencies relative to that of native MLucV were calculated as follows:

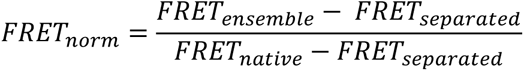

where FRET_ensemble_ is the measured ensemble FRET efficiency, FRET_separated_ is the calculated ensemble FRET measured in a solution of separated mTFP1 and Venus (0.33) and FRET_native_ is the measured ensemble FRET of native MLucV (0.43). Unless otherwise specified, all ensemble FRET measurements were performed at 400 nM of MLucV. The temperature was maintained at 25°C unless otherwise specified. All experiments were performed in LRB (20mM Hepes-KOH pH7.4,150mM KCl, 10mM MgCl2) refolding buffer containing 5 mM ATP, 2mM DTT, unless otherwise specified. 4μM BSA was used in assays with chaperones to avoid MLucV species sticking to the vessel, it does not affect the fate of the formed aggregates, nor affects the activity of the chaperones. All experiments were repeated at least three times.

### Analysis of proteins models

All the models (PDBs) have been analyzed and rendered with UCSF ChimeraX tool (Meng, Goddard et al. 2023).

## Funding

Swiss National Fund grant 31003A_175453

## Author contributions

Conceptualization: MER, PG, PDLR

Methodology: PG, PDLR, MER, BF, ST, AM

Investigation: PG, PDLR, MER, BF, ST, AM

Visualization: PG, PDLR, MER, BF, ST, AM

Supervision: PG, PDLR

Writing—original draft: PG, PDLR, MER

Writing—review & editing: PG, PDLR, MER, BF, ST

## Data and materials availability

All data are available in the main text or supplementary materials.

## Competing interests

The authors declare that they have no competing interests.

## Supplementary Data

Supplementary Information: Sequences of Wildtypes proteins Sis1 (SS) and Ydj1(YY) and both SWAPS. Sis1’ JD is highlighted in cyan, and Ydj1’a JD in green.

>SS

MVKETKLYDLLGVSPSANEQELKKGYRKAALKYHPDKPTGDTEKFKEISEAFEILNDPQKREIYDQYGLEAARSGGPSFGPGGPGGAGGAGGFPGGAGGFSGGHAFSNEDAFNIFSQFFGGSSPFGGADDSGFSFSSYPSGGGAGMGGMPGGMGGMHGGMGGMPGGFRSASSSPTYPEEETVQVNLPVSLEDLFVGKKKSFKIGRKGPHGASEKTQIDIQLKPGWKAGTKITYKNQGDYNPQTGRRKTLQFVIQEKSHPNFKRDGDDLIYTLPLSFKESLLGFSKTIQTIDGRTLPLSRVQPVQPSQTSTYPGQGMPTPKNPSQRGNLIVKYKVDYPISLNDAQKRAIDENF

>SY

MVKETKLYDLLGVSPSANEQELKKGYRKAALKYHPDKPTGDTEKFKEISEAFEILNDPQKREIYDQYGLEAARSGGPSFGGGFPGGGFGFGDDIFSQFFGAGGAQRPRGPQRGKDIKHEISASLEELYKGRTAKLALNKQILCKECEGRGGKKGAVKKCTSCNGQGIKFVTRQMGPMIQRFQTECDVCHGTGDIIDPKDRCKSCNGKKVENERKILEVHVEPGMKDGQRIVFKGEADQAPDVIPGDVVFIVSERPHKSFKRDGDDLVYEAEIDLLTAIAGGEFALEHVSGDWLKVGIVPGEVIAPGMRKVIEGKGMPIPKYGGYGNLIIKFTIKFPENHFTSEENLKKLEEILPPRIVPAIPKKATVDECVLADFDPAKYNRTRASRGGANYDSDEEEQGGEGVQCASQ

>YY

MVKETKFYDILGVPVTATDVEIKKAYRKCALKYHPDKNPSEEAAEKFKEASAAYEILSDPEKRDIYDQFGEDGLSGAGGAGGFPGGGFGFGDDIFSQFFGAGGAQRPRGPQRGKDIKHEISASLEELYKGRTAKLALNKQILCKECEGRGGKKGAVKKCTSCNGQGIKFVTRQMGPMIQRFQTECDVCHGTGDIIDPKDRCKSCNGKKVENERKILEVHVEPGMKDGQRIVFKGEADQAPDVIPGDVVFIVSERPHKSFKRDGDDLVYEAEIDLLTAIAGGEFALEHVSGDWLKVGIVPGEVIAPGMRKVIEGKGMPIPKYGGYGNLIIKFTIKFPENHFTSEENLKKLEEILPPRIVPAIPKKATVDECVLADFDPAKYNRTRASRGGANYDSDEEEQGGEGVQCASQ

>YS

MVKETKFYDILGVPVTATDVEIKKAYRKCALKYHPDKNPSEEAAEKFKEASAAYEILSDPEKRDIYDQFGEDGLSGAGGAPGGPGGAGGAGGFPGGAGGFSGGHAFSNEDAFNIFSQFFGGSSPFGGADDSGFSFSSYPSGGGAGMGGMPGGMGGMHGGMGGMPGGFRSASSSPTYPEEETVQVNLPVSLEDLFVGKKKSFKIGRKGPHGASEKTQIDIQLKPGWKAGTKITYKNQGDYNPQTGRRKTLQFVIQEKSHPNFKRDGDDLIYTLPLSFKESLLGFSKTIQTIDGRTLPLSRVQPVQPSQTSTYPGQGMPTPKNPSQRGNLIVKYKVDYPISLNDAQKRAIDENF

---

**Figure S1:**
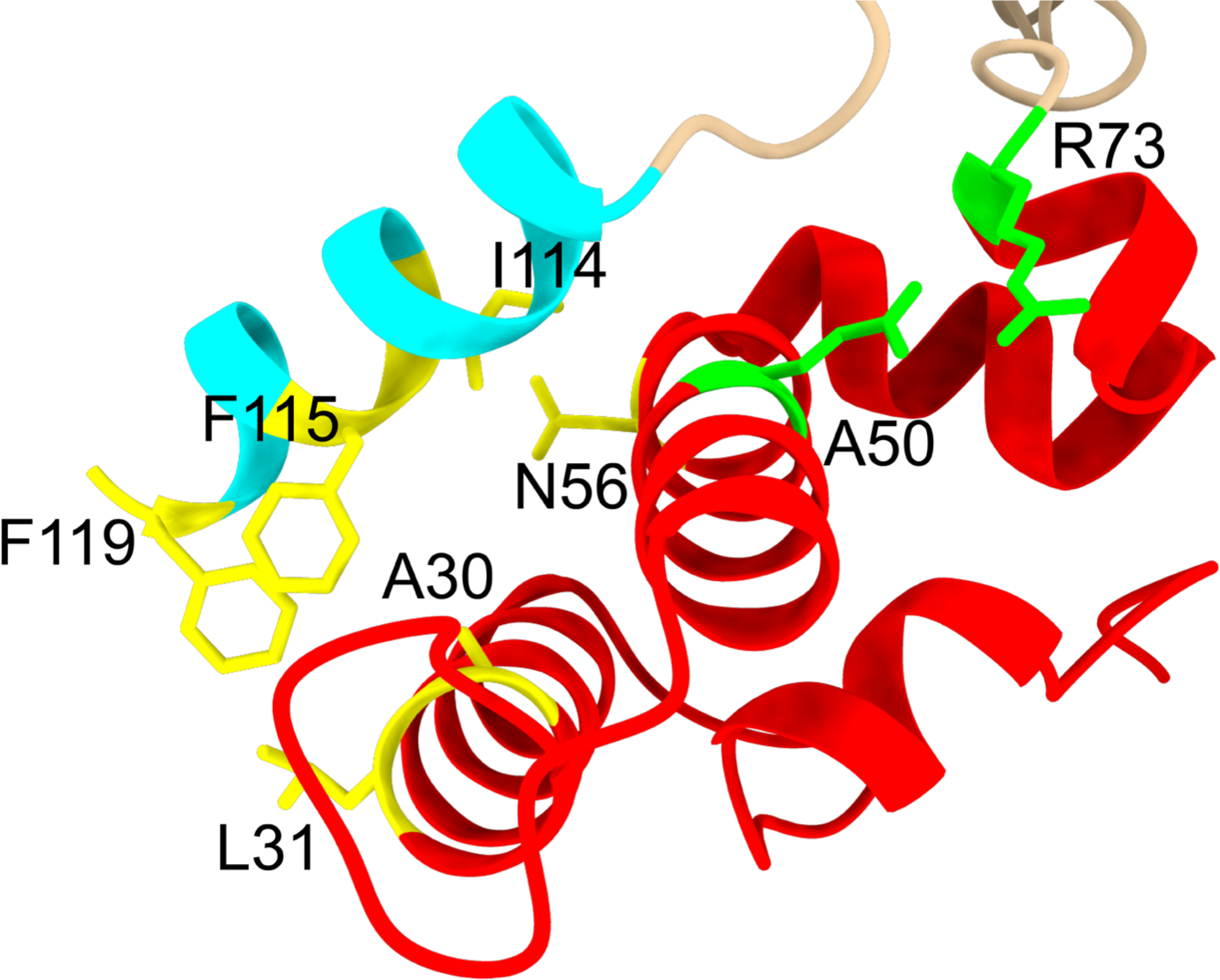
Blow-up on the J-domain and Helix V of Alphafold generated structure of SIS-1. The interaction between Glu-50 on JD (red) and Arg-73 (green) on a short helical motif immediately C-terminal to helix IV and the Helix V (blue) proximity with the JD are modelled here based on the Alphafold prediction of Sis1 (Uniprot ID: AF-P25294-F1). For the interactions between helices II and III with V (yellow), the contacts have been derived from the aligned sequence of Sis1 with the one of DNAJB1 from the work of Faust (Faust, Abayev-Avraham et al. 2020). For clarity, most of the surrounding model of Sis1 has been hidden.

**Figure S2:**
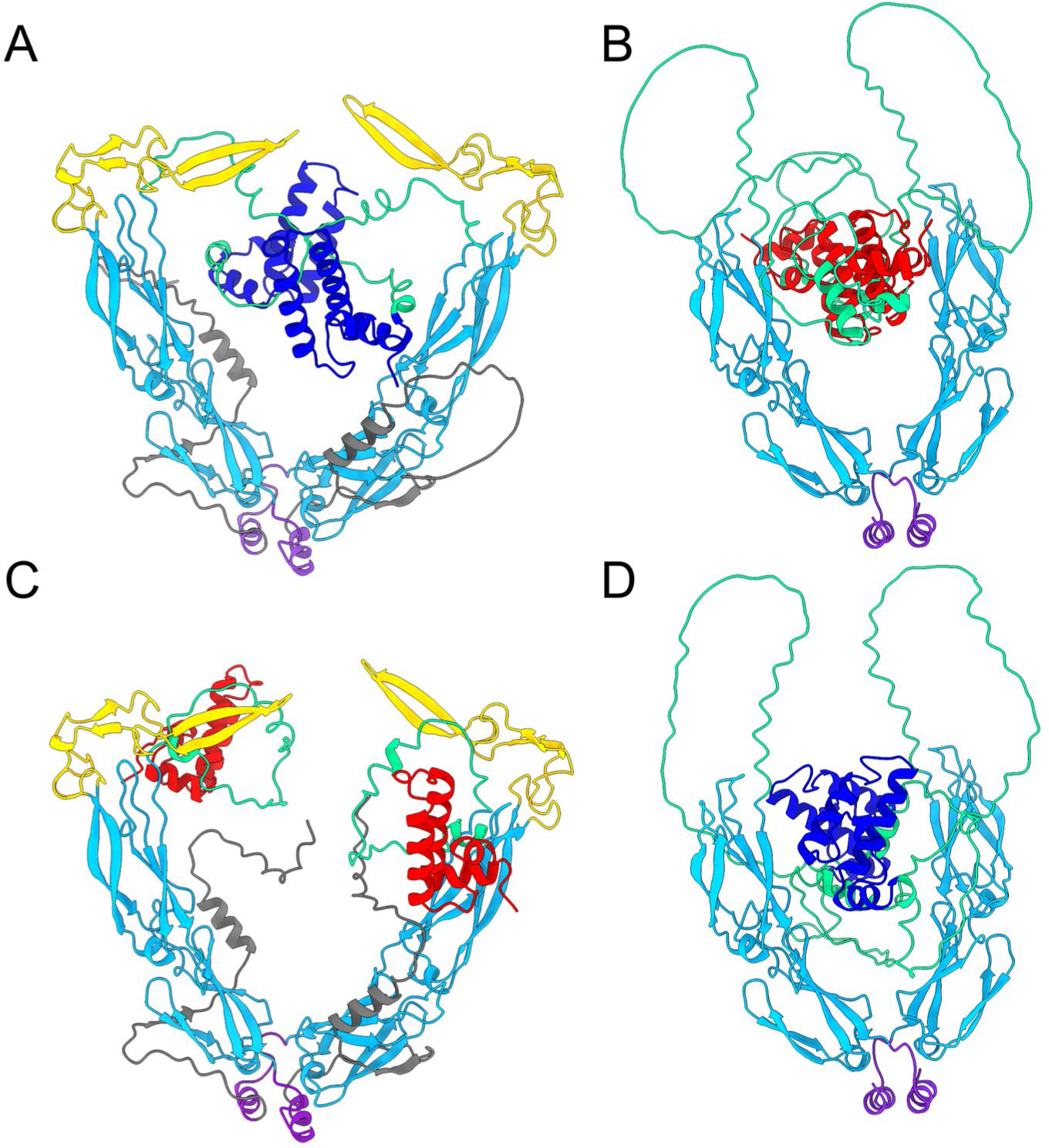
Alphafold 2 predicted structures of the different JDPs. A: YY, B: SS, C: SY: D: YS

**Figure S3:**
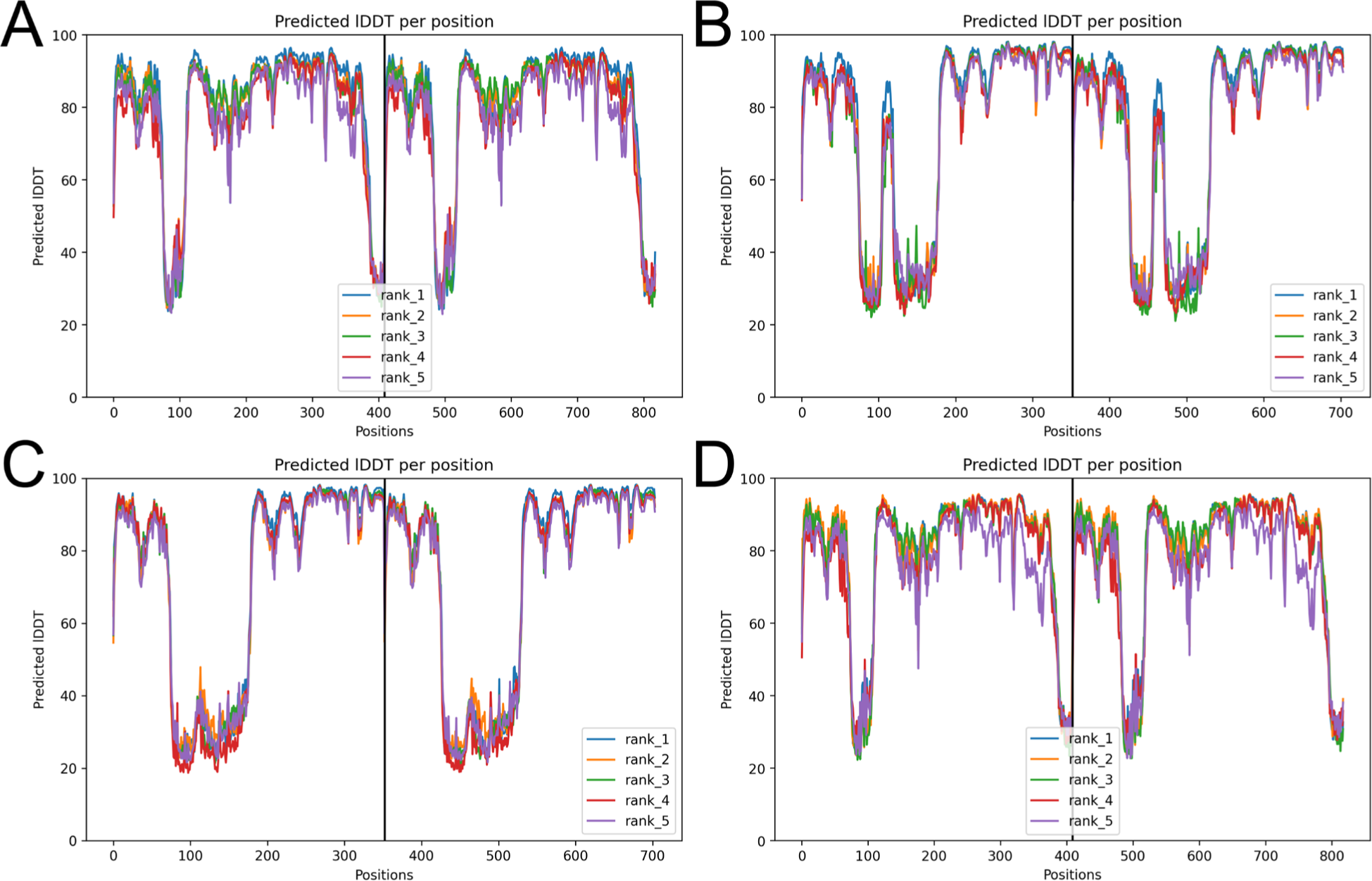
PLDDT values of the 4 JDPs from Alphafold models in Figure S1 for A: YY, B: SS, C: SY: D: YS.

**Figure S4:**
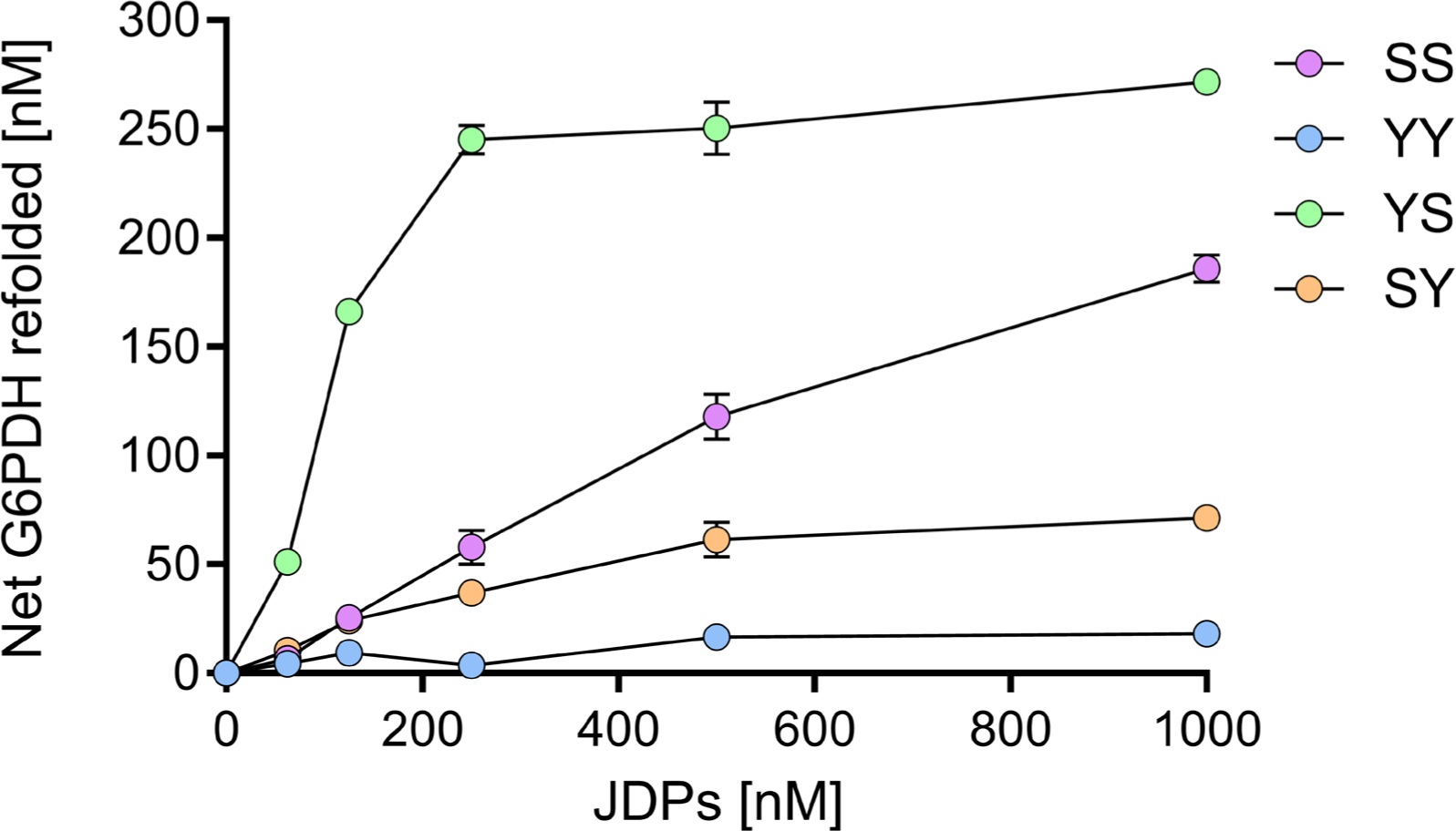
Efficiency of four different JDPs proteins to activate the SSA1-mediated disaggregation of heat-denatured G6PDH. 0.5 µM of stable inactive G6PDH aggregates were incubated up to 120 minutes at 25°C in the presence of 5 mM ATP, 6 µM of SSA1, 0.75 µM SSE1, and between 0 to 1 µM of either YY, SS, SY or YS. G6PDH activity was measured at different times of chaperone-mediated refolding reaction at 25°C. In all panels, error bars represent mean ± SD (n = 3).

**Figure S5:**
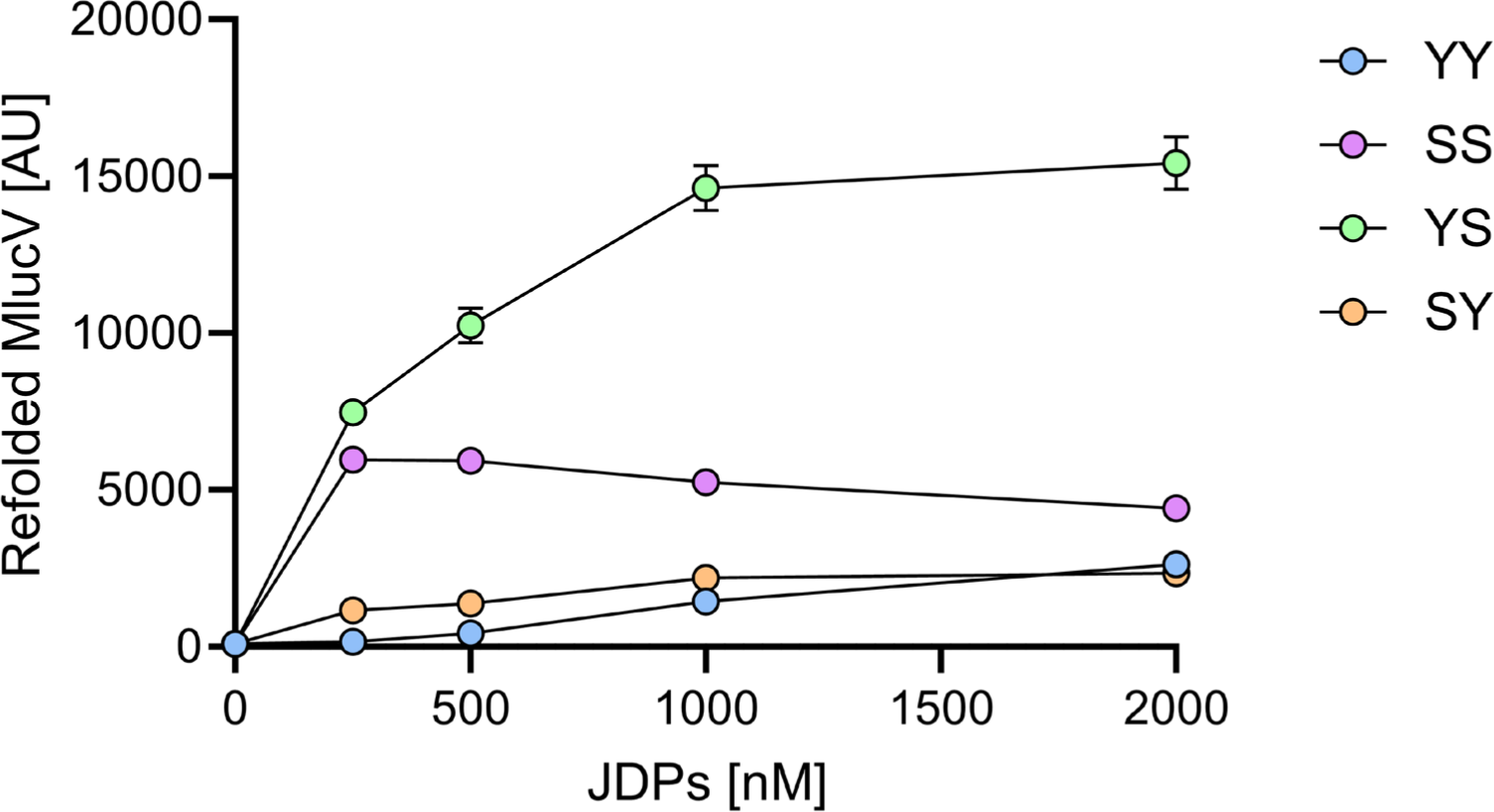
MLucV Refolding assay. Stable urea-inactivated MLucV aggregates (Tiwari, Fauvet et al. 2023) were incubated up to 120 minutes at 25°C in the presence of 5 mM ATP, 6 µM of SSA1, 0.75 µM SSE1, and between 0 to 2 µM of either YY, SS, SY or YS. In all panels, error bars represent mean ± SD (n = 3).

**Figure S6:**
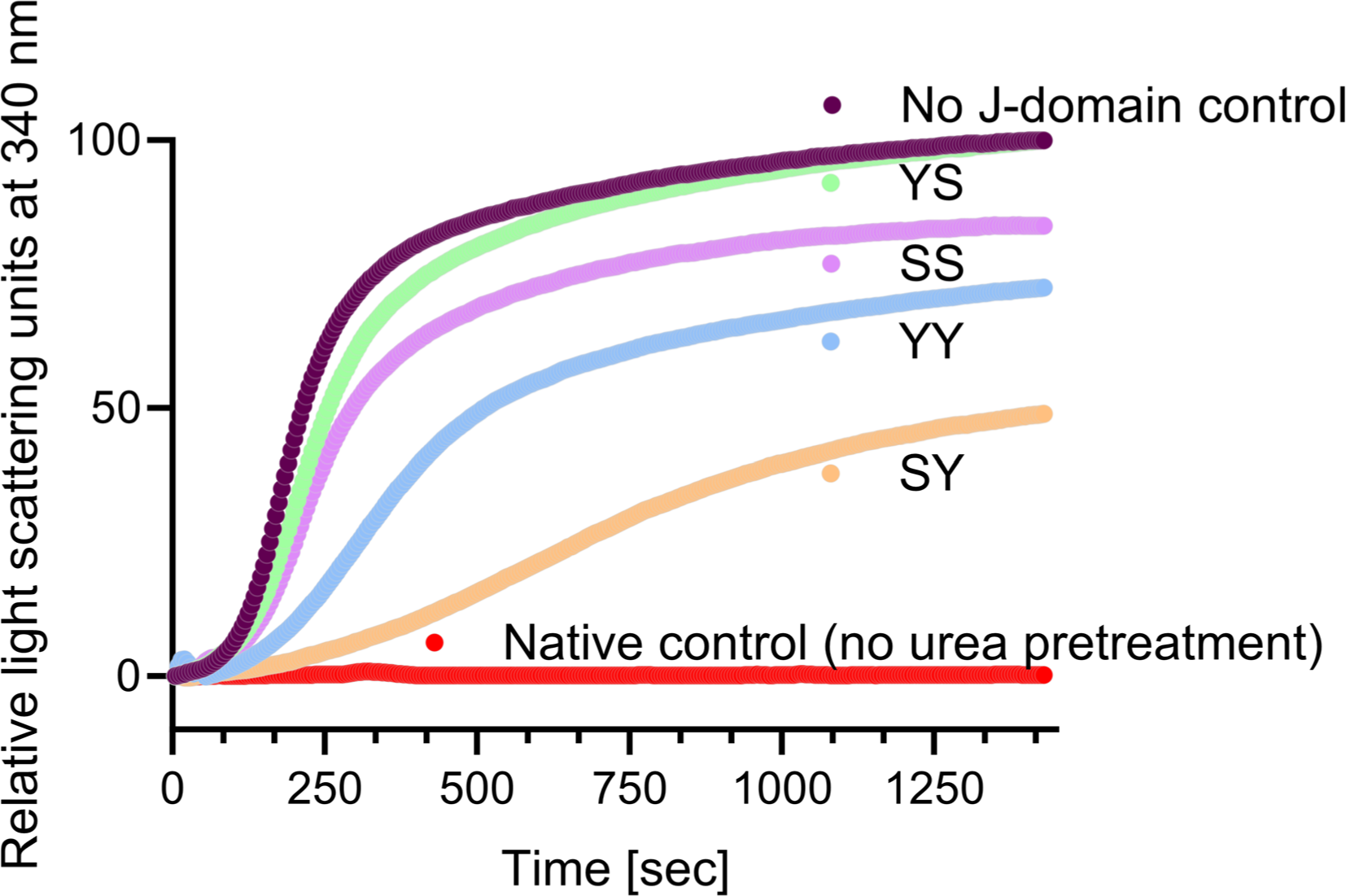
J-domain protein acting as holding chaperone10 µM Luciferase was denatured in 6M Urea at 30℃ for 30 min, then diluted to 0.3µM in 50 mM Tris-pH 7.5+150mM KCl+10mM Mgcl2+2mM DTT buffer in the absence or in the presence of 2µM of YY, SS, SY, YS. The time –dependent aggregation was monitored by light scattering at 340 nm, at 30℃. . 100% of aggregation was set on the urea denatured Luciferase without any addition of JDPs.

**Figure S7:**
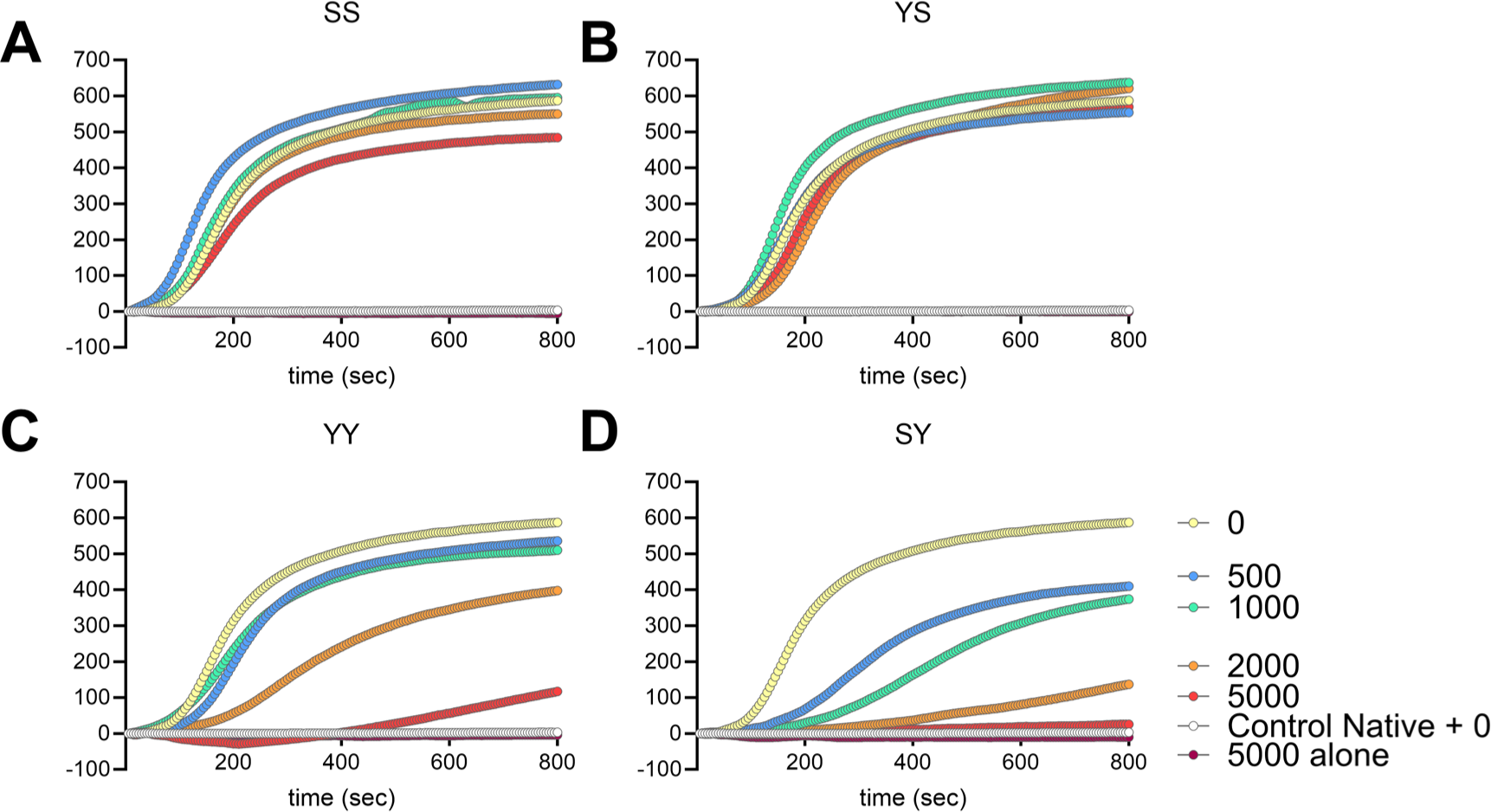
Prevention of aggregation of urea-pre-unfolded luciferase by increased YY, SS, SY, and YS concentrations. 10 µM luciferase was preincubated in 6 M urea at 30℃, then diluted to a final concentration of 0.3 µM in buffer containing 0, 0.5, 1, 2, 5 µM of SS (A), YS (B), YY (C), SY (D). Aggregation was monitored by light scattering at 340 nm at 30°C.

**Figure S8:**
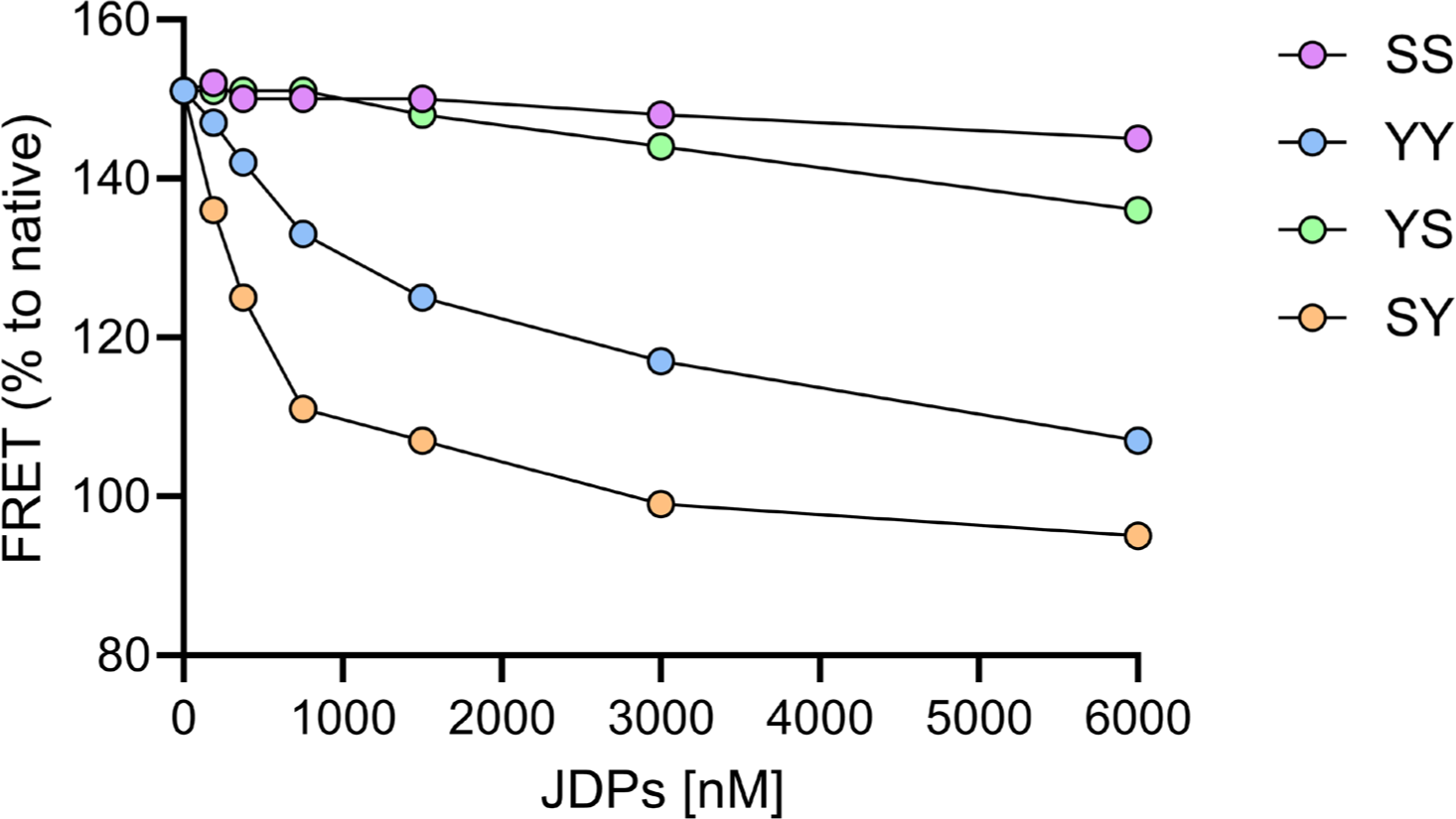
200 nM of native MlucV (Tiwari, Fauvet et al. 2023) in the presence of 4µM BSA and indicated concentrations (0, 187.5, 375, 750, 1500, 3000, 6000 nM) of JDPs were incubated for 360 seconds at 38°C, following which, only 0.1% of the luciferase was still active. Luciferase activity and FRET efficiency were measured at the indicated time points and expressed as % of the initial luciferase activity and FRET efficiency at t=0. Because time-dependent measures of protein aggregation by light scattering are notoriously variable the presented results are only representative of three replicates.

**Figure S9:**
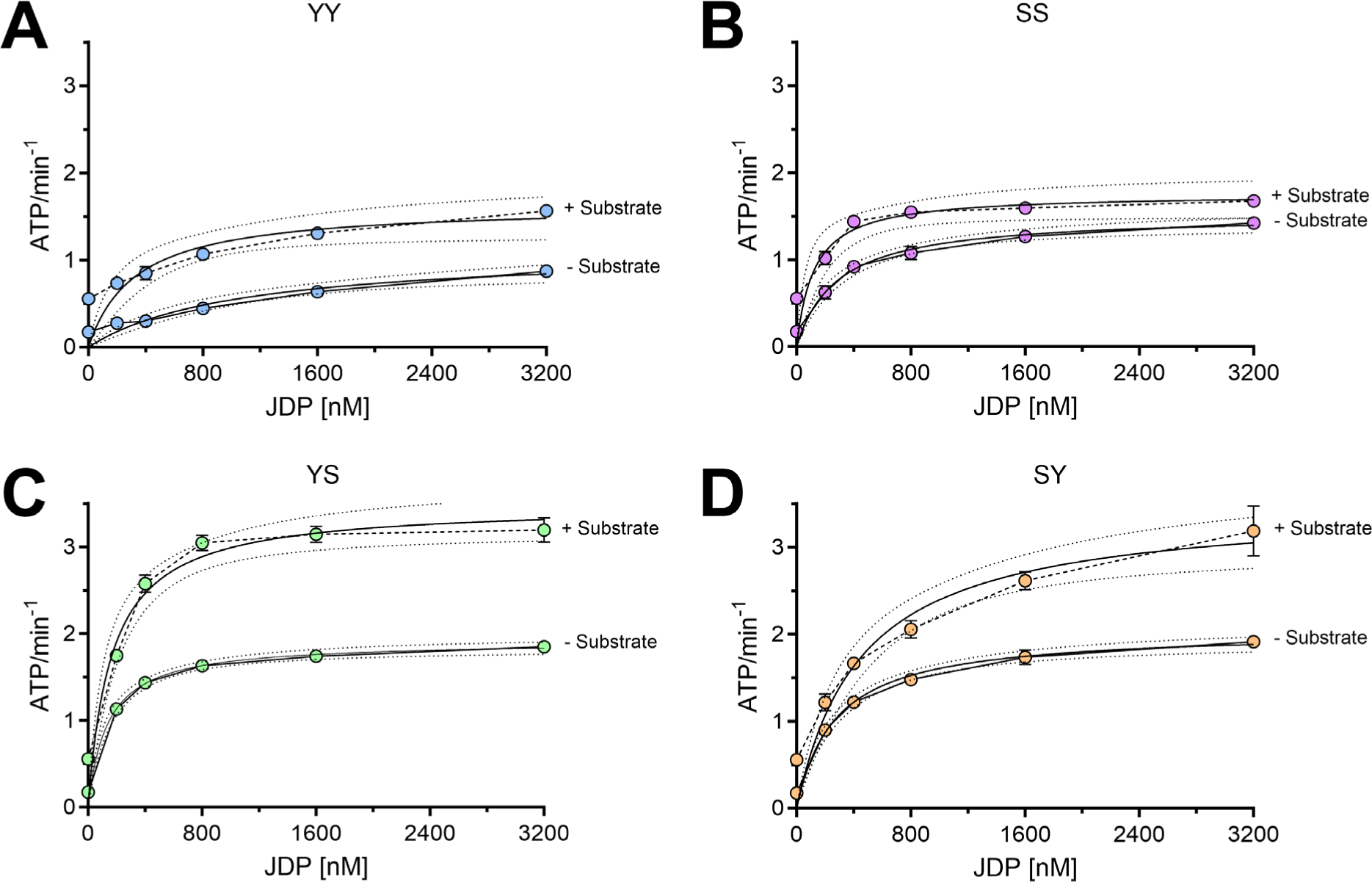
Substrate and JDPs increase ATPase activity of Hsp70 (SSA1). A-D: Rates of ATP hydrolysis of 6000 nM SSA1, and 750 nM SSE1 in the presence of increasing concentrations of YY, SS, YS, and SY, with or without 200 nM of pre-aggregated luciferase. The error bars in all panels represent the SD between replicates (n = 3).

**Figure S10:**
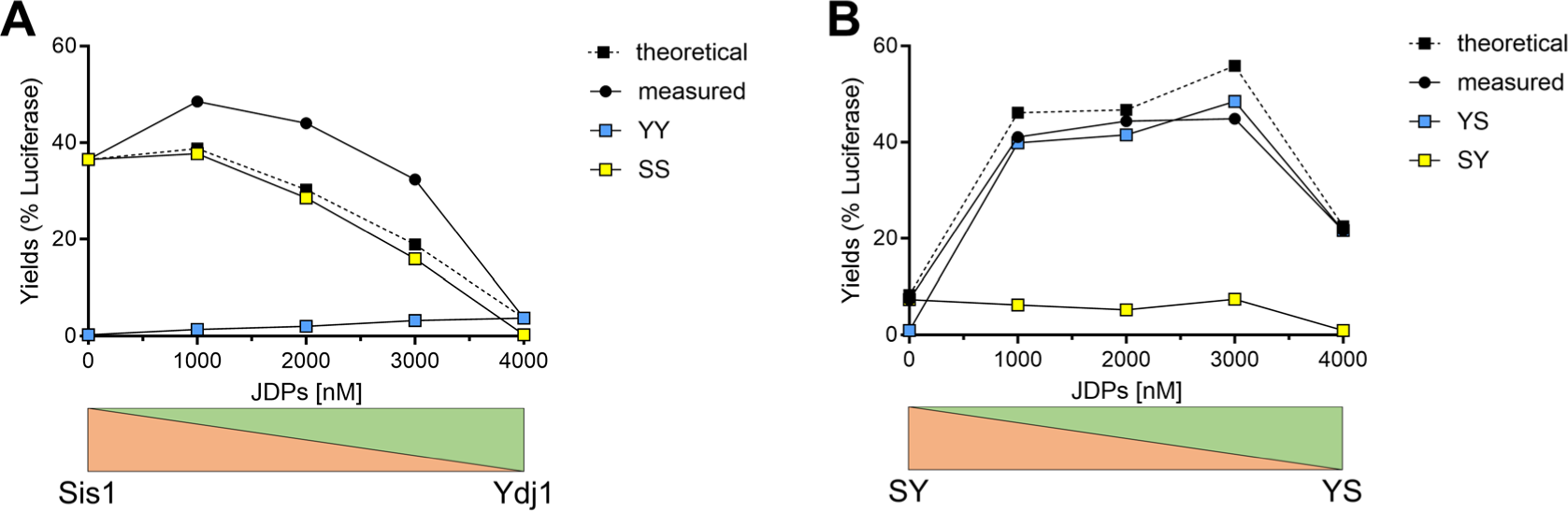
Mixture of YY and SS and YS and SY and refolding yields. A: Refolding yields of heat-pre-aggregated luciferase (200 nM) in the presence of SSA1 (6 µM), SSE1 (0.75 µM), ATP (4 mM) at indicated SS: YY ratios, between 4 µM SS and 0 µM YY (left), up to 4µM YY and 0 µM SS (right). Hatched lines, theoretical values if SS does not collaborate with YY. B: Refolding yields of heat-pre-aggregated luciferase (200 nM) in the presence of SSA1 (6 µM), SSE1 (0.75 µM), ATP (4 mM) at indicated SY: YS ratios, between 4 µM SY and 0 µM YS (left), up to 4 µM YS and 0 µM SY (right). Hatched lines, theoretical values if SY does not collaborate with YS.

## Notes

### Competing Interest Statement

The authors have declared no competing interest.

